# Migration through a small pore disrupts inactive chromatin organization in neutrophil-like cells

**DOI:** 10.1101/339085

**Authors:** Elsie C. Jacobson, Jo K. Perry, David S. Long, Ada L. Olins, Donald E. Olins, Bryon E. Wright, Mark H. Vickers, Justin M. O’Sullivan

## Abstract

**Background:** Mammalian cells are flexible and can rapidly change shape when they contract, adhere, or migrate. Their nucleus must be stiff enough to withstand cytoskeletal forces, but flexible enough to remodel as the cell changes shape. This is particularly important for cells migrating through constricted space, where the nuclear shape must change in order to fit through the constriction. This occurs many times in the life cycle of a neutrophil, which must protect its chromatin from damage and disruption associated with migration.

**Results:** Total RNA-sequencing identified that neutrophil migration through 5 or 14μm pores was associated with changes in the transcript levels of inflammation and chemotaxis-related genes, when compared to unmigrated cells. Differentially expressed transcripts specific to migration with constriction were enriched for groups of genes associated with cytoskeletal remodeling.

Hi-C was used to capture the genome organization in control and migrated cells. Minimal switching was observed between the active (A) and inactive (B) compartments after migration. However, global depletion of short range contacts was observed following migration with constriction compared to migration without constriction. Regions with disrupted contacts, TADs, and compartments were enriched for inactive chromatin.

**Conclusion:** Short range genome organization is preferentially altered in inactive chromatin, possibly protecting transcriptionally active contacts from the disruptive effects of migration with constriction. This is consistent with current hypotheses implicating heterochromatin as the mechanoresponsive form of chromatin. Further investigation concerning the contribution of heterochromatin to stiffness, flexibility, and protection of nuclear function will be important for understanding cell migration in human health and disease.

## Background

Mammalian cells are subject to a wide range of mechanical environments and processes, including fluid shear stress [1], matrix stiffness [2], and migration [3]. Mounting an appropriate response to these stimuli is crucial for blood vessel development [1], cardiac health [4], muscle tone [5], bone strength [6], and the immune response [7]. Dysfunctional responses can lead to a wide range of diseases [8].

Cells are mechanically connected to their surroundings by integrins and other mechanoresponsive cell surface proteins [9]. In turn, these mechanoresponsive cell surface proteins are connected to the cytoskeleton, which remodels in response to chemical and mechanical signalling (reviewed in [10]). Cytoskeletal remodeling controls the shape of the cell, the nucleus, and even chromatin organization [11–13]. External mechanical signals pass directly to the nucleoskeleton [14] through the linker of nucleoskeleton and cytoskeleton (LINC) complex, which connects the cytoskeleton to the nucleus [15]. A major mechanoresponsive element of the nucleoskeleton is the nuclear lamina, a key structural element which forms a mesh around the inside of the nuclear envelope [16]. The mammalian lamina consists of two major forms of intermediate filament; A-type lamins (Lamin A and C) and B-type lamins (Lamin B1, B2) [17]. High levels of Lamin A/C result in a stiffer nucleus [2,18,19], although it is becoming increasingly recognized that chromatin also contributes to nuclear mechanics [20,21]. Heterochromatin stiffens the nucleus [20], interacts preferentially with the nuclear lamina, [22] the LINC complex [23], and other nuclear envelope proteins [24] and is therefore believed to play a role in the mechanoresponsiveness of the nucleus [20,25].

There are many acute consequences of mechanically challenging the nucleus [26,27]. When endothelial cell nuclei are exposed to shear stress, the position and shape of the nuclei aligns with the direction of the shear stress [1]. This alignment extends to loci within the nucleus, which move in the direction of the force after exposure to shear stress for more than 30 minutes [28]. Shear or stiffness induces cytoplasmic to nuclear shuttling of transcriptional and chromatin remodeling proteins including yes-associated protein (YAP)/transcription coactivator with PDZ binding domain (TAZ) [29] and chromatin modifier histone deacetylase 3 (HDAC3) [12]. In response to acute, applied force, chromatin linearizes [30,31] and becomes more transcriptionally active [31]. However nuclear remodeling that occurs during cell migration is associated with chromatin condensation, with chemically-induced decondensation reducing the rate of migration [25,32].

Migration through a constriction provides unique mechanical challenges [33]. Due to its size and stiffness, the nucleus is the rate limiting step for size reduction [18,34], but it also provides a stiff internal surface for the cytoskeleton to ‘push’ on and generate force [35]. Nuclear positioning during migration is tightly controlled by the cytoskeleton, and it is usually positioned towards the back of the cell [36]. However, the nucleus leads transendothelial migration in immune cells, with nuclear lobes ‘drilling’ between endothelial cells to increase the gap between them and allow the neutrophil to pass through [37,38].

The nucleus plays a critical structural role during migration through constricted space, yet the spatial organization of DNA is critical for transcriptional regulation, [39] and neutrophils must retain the structural features that are required for normal gene expression. Heterochromatin is tightly packed, self-interacting, and tethered at the periphery of most cells, while transcriptionally active euchromatin is found towards the center [40]. Within these compartments, local clusters of DNA are formed by loop extrusion [41,42], increasing the interactions between enhancers and promoters that are found within each topologically associated domain (TAD) [43]. Beyond individual enhancer/promoter interactions [44], active genes and enhancers cluster together [45] around the transcriptional machinery [46]. While many studies have described chromatin reorganization during differentiation and disease (for example, [47–49]), it is not known how genome organization is affected by the acute shape changes that occur during migration with constriction. Neutrophils are highly migratory cells, with features that may contribute to nuclear stability, including a lobed nucleus, low Lamin A, and high heterochromatin content [18,25,50].

Here, we investigated whether migration with constriction has an acute impact on chromatin conformation and transcriptional activity using an *in vitro* cell migration model. Neutrophil-like HL-60/S4 cells were differentiated and migrated through 5μm pores (constricted migration) or 14μm pores (non-constricted migration). When the two migration conditions were compared to each other, and to unmigrated cells, RNA-seq identified a strong transcriptional response to migration, and additional effects associated with constriction. Hi-C analysis demonstrates that global genome structure is largely maintained following constricted migration, with local disruptive effects occurring preferentially in inactive chromatin.

## Results

### An in vitro migration model

To investigate the effects of acute cell shape change, we developed a simplified model of neutrophil migration. HL-60/S4 cells were differentiated into neutrophil-like cells using retinoic acid (HL-60/S4-RA) [51]. Increased cell surface CD11b and nuclear lobulation confirmed differentiation (Additional file 1: Supplementary fig 1a and b). Cell viability was confirmed with flow cytometric analysis of annexin V immunolabelled and propidium iodide stained cells (Additional file 1: Supplementary fig 1c). HL-60/S4-RA cells were migrated through two pore sizes, 5μm and 14μm, to simulate migration with and without constriction (Fig 1a).

**Figure 1.**
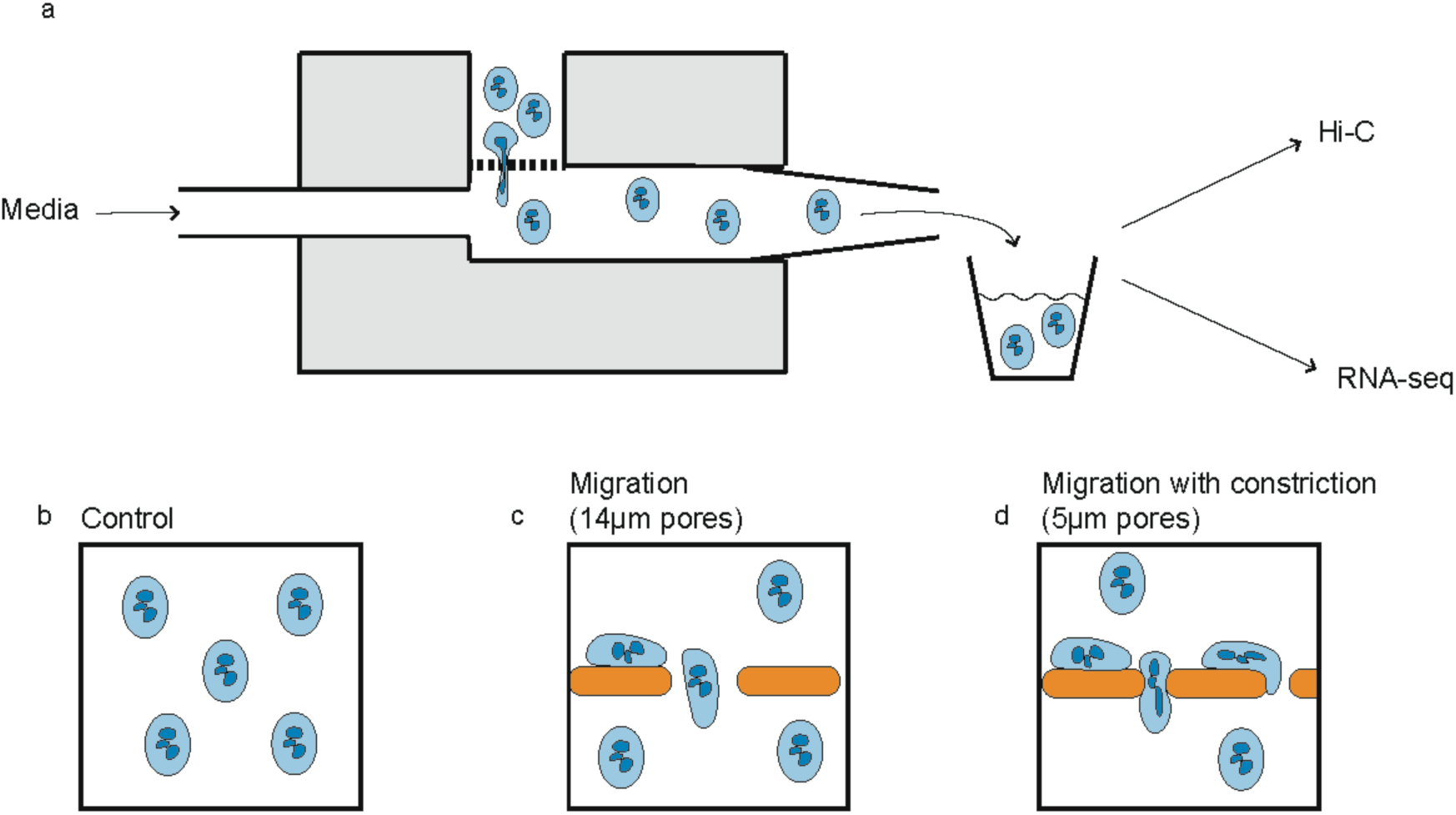
Neutrophil-like cells were migrated through two sizes of porous membrane. a) A classical Boyden chamber was modified to allow a continuous flow of media (0.2 mL/min) through the lower well. Migrated cells were processed every thirty minutes. Three experimental conditions were used. b) Cells that did not migrate c) cells that migrated through pores larger than themselves (14μm diameter) and d) cells that migrated through pores smaller than themselves (5μm diameter).

Consistent with previous reports [18], viable HL-60/S4-RA cells had an average diameter of 7.0 ± 0.3μm (Additional file 1: Supplementary fig 1d) and passed through the 14μm pores rapidly. By contrast, the rate of migration through 5μm pores was much slower, most likely due to the remodeling required for a cell to fit through a pore smaller than itself (Additional file 1: Supplementary fig 1e). Migration rate was unlikely to be affected by pore availability as electron microscopy confirmed that the 5 and 14μm pores cover approximately 9% or 5% of the membrane surface, respectively (Additional file 1: Supplementary fig 2). No additional chemoattractants were added to the lower well, as the gradients of both cell density and serum were sufficient to induce migration. The small pores used in our assay do not require extreme deformation. However, increased nuclear stiffness significantly slows migration rate of HL-60/S4-RA cells through 5μm diameter pores, consistent with nuclear remodeling representing the rate-limiting step in this assay [18].

**Figure 2.**
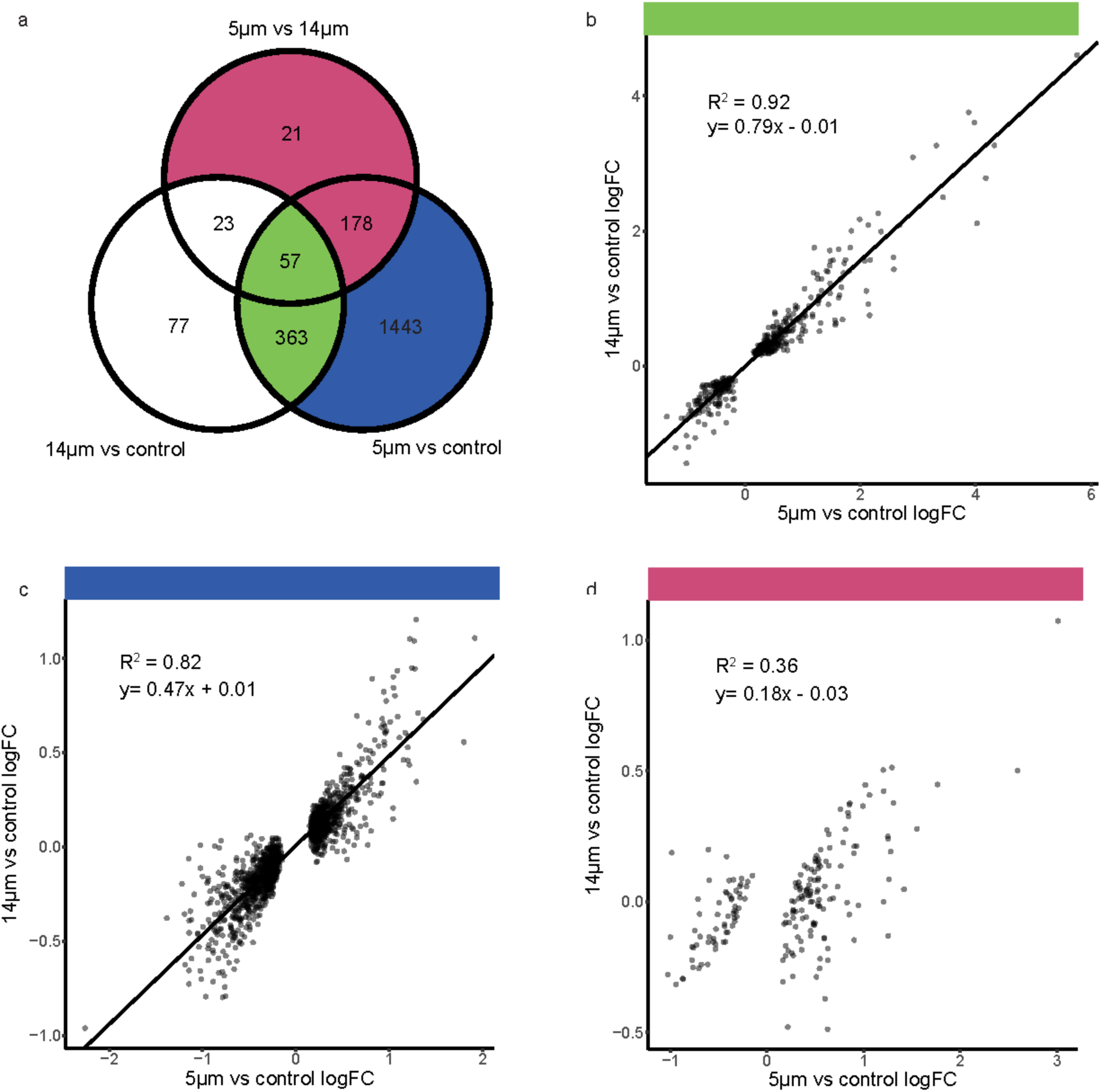
Gene expression changes. a) Venn diagram of significantly differentially expressed genes (DESeq2, FDR<0.05) between 14μm and 5μm pores, unmigrated and 5μm pores, unmigrated and 14μm pores. 80% of the genes differentially expressed after migration through 14μm pores also change after migration through 5μm pores. b) The 420 genes that changed in both conditions had highly correlated (R^2^=0.92) log2 fold changes (logFC). c) While 1621 additional genes change expression after migration through 5μm, only 178 of these were significantly different between the two pore sizes. The 1443 that were not significantly different had highly correlated changes compared to unmigrated cells (R^2^=0.82). d) The relatively small subset of 199 genes are uniquely different after migration through 5μm pores. We believe these genes are associated with the effects of remodeling and have low correlation (R^2^=0.36) between 5 and 14μm pore migration.

### Transcriptional changes after migration and remodeling are distinct

We captured acute responses to migration with or without constriction by processing cells within 30 minutes post-migration, when early responses to inflammatory stimuli can be detected [52]. We contend that this model allows us to isolate the effects of passage through the transwell chamber from the effects of nuclear remodeling that resulted from constricted migration. We used total RNA-seq to identify genes that were differentially expressed, comparing unmigrated control cells with cells that migrated through 5μm or 14μm pores. This enabled us to identify changes in steady-state total RNA levels associated with migration per se, and to distinguish them from transcriptional changes specific to migration with constriction.

Comparisons of gene transcript levels between cells migrated without constriction and unmigrated cells identified 304 genes that were significantly upregulated and 216 significantly downregulated genes (FDR<0.05). Comparisons of gene transcript levels between cells migrated with constriction and unmigrated cells identified 1057 significantly upregulated and 984 significantly downregulated genes (FDR<0.05). Comparisons of cells migrated with and without constriction identified 204 significantly upregulated and 75 significantly downregulated genes (FDR<0.05) (Additional file 1: Supplementary fig 3b,c,d; Additional file 2: Supplementary table 1).

**Figure 3.**
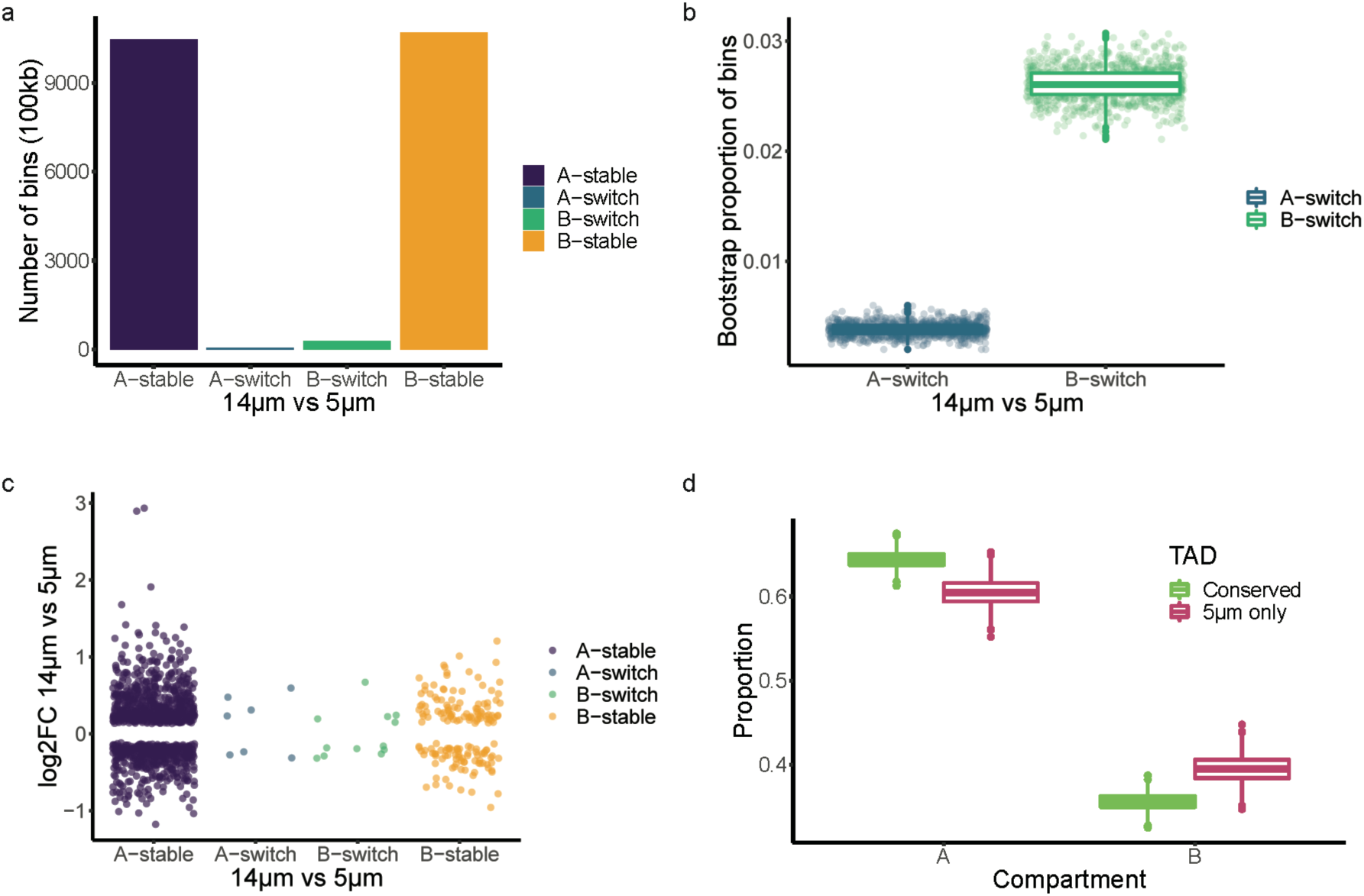
Compartment switching following migration with or without constriction. a) Compartment structure is mostly conserved, with a small proportion of regions in different compartments in cells migrated with or without constriction. 100kb bins were considered switched if they had the opposite PC1 sign and low correlation (R<0.6) between the two conditions. b) Bootstrapping found that switching from compartment B occurred more frequently than switching from compartment A (p<2.2×10^−16^, logistic regression). c) Compartment switching was not associated with gene expression changes. The log2 fold change of differentially expressed genes (5μm vs 14μm pores) did not correlate with compartment or switching status. d) TADs identified in migration with constriction were considered conserved if they overlapped <80% with TADs identified in migration without constriction. Conserved TADs were more likely to be found in compartment A than non-conserved TADs (ANOVA with Tukey adjustment, adjusted p<3×10^−8^).

The majority of observed migration-associated gene expression changes fell into one of three sets. The first set included 420 genes with highly correlated fold changes following migration through 5μm or 14μm pores, when compared to unmigrated control cells (R^2^=0.92, p < 2.2e-16, Fig 2a and b). We consider this set of genes to reflect the effects of passage through the migration chamber, which includes changing cell density, exposure to fresh media, and fluid shear. Gene ontology (GO) analysis, using TopGo [53], identified enrichment for GO biological process terms that included response to cytokine (FDR adjusted p = 4.06e^-05^, Fisher’s exact test) and monocyte chemotaxis (FDR adjusted p = 0.044, Fisher’s exact test; Additional file 3: Supplementary table 2). This suggested that the environment of the migration chamber was sufficient to stimulate immune-like transcriptional responses in these neutrophil-like cells, despite the lack of added chemoattractants, and 14μm diameter pores being substantially larger than the 7μm diameter cells [18] (Additional file 1: Supplementary fig 1d).

The second set contained 1433 genes that were significantly differentially expressed after migration through 5μm pores, when compared to unmigrated control cells. The genes within this group were not significantly differentially expressed following migration through 14μm pores. However, the highly correlated fold changes (R^2^=0.82, p < 2.2e^-16^) indicated that the change in RNA transcript levels following migration without constriction was in the same direction, but with larger effect sizes, following migration with constriction (Fig 2a and c). GO analysis identified enrichment for terms, including inflammatory response (FDR adjusted p = 0.004, Fisher’s exact test) and chemokine mediated signalling pathway (FDR adjusted p = 0.011, Fisher’s exact test) (Additional file 3: Supplementary table 3).

The third set of 199 genes exhibited differential transcript levels following migration through 5μm pores when compared to either 14μm pores or control cells. There was a low correlation between the transcript levels of these genes following migration through the two pore sizes, compared to unmigrated cells (R^2^=0.32, p < 2.2e^-16^, Fig 2a and d). Therefore, this set represents the transcript changes that were specific to the constriction aspect of the assay. GO analysis demonstrated that genes associated with distinct biological processes were regulated in the cells migrated with constriction. Seventeen genes were involved with actin cytoskeleton remodeling (FDR adjusted p = 0.071, Fisher’s exact test) (Additional file 3: Supplementary table 4). In addition, migration through 5μm pores was associated with changes in transcript levels of genes with molecular function associated with lactate transport (FDR adjusted p = 0.025, Fisher’s exact test), and Rab GTPase activity (FDR adjusted p = 0.086, Fisher’s exact test) (Additional file 3: Supplementary table 5).

Neutrophil-like cells are more migratory than their promyelocytic precursors [54]. Therefore, it was possible that migration might select for cells with a stronger neutrophil-like phenotype. To determine whether migration had selected for neutrophil-like cells we compared gene expression changes of cells migrated with and without constriction, with changes that occurred during differentiation into neutrophil-like cells [55]. There was no obvious relationship in the gene expression profiles across these two groups (Additional file 1: Supplementary fig 3e). Therefore our *in vitro* migration model did not obviously select for a subpopulation of highly neutrophilic cells.

Collectively, the results of the total RNA-seq were consistent with migration through either pore size being associated with changes in transcript levels for genes associated with inflammation and chemotaxis. Moreover, migration with constriction was associated with changes in transcript levels of cytoskeleton remodeling genes.

### Features of genome organization are more stable in compartment A

We used Hi-C to investigate global features of genome organization, to determine whether migration induced remodeling at different levels of nuclear structure. Heterochromatin preferentially interacts with itself, forming a phase separated compartment isolated from euchromatin [40,56]. Distinct interaction patterns can be extracted from the Hi-C heatmap using principle component analysis, allowing us to assign regions of DNA to either the active (A) or inactive (B) compartment, which generally corresponds to euchromatin and heterochromatin respectively [57]. PC1 values for 100kb bins were calculated using HOMER [58]. Bins with positive PC1 values were assigned to compartment A, while bins with a negative PC1 value were assigned to compartment B. Correlation matrices of migrated and unmigrated cells were compared directly in order to identify regions that showed different interaction patterns between conditions. 99.6% of A compartment bins and 97.4% of B compartment bins had the same compartment assignment and a correlation >0.6 and were thus considered stable between migration with and without constriction (Fig 3a). Although compartment structure was generally conserved, there was significantly more disruption to the B compartment (logistic regression, p<2×10^−16^) (Fig 3b).

If altered gene expression was simply a result of genes switching between compartments, we would expect to see increased RNA expression for genes that moved from compartment B to A, and decreased expression for those moving from A to B. However, there was no correlation between the direction of change of differentially expressed genes and region stability (ANOVA with Tukey adjustment, p>0.6 for all comparisons) (Fig 3c).

TADs are conserved units of DNA organization characterized by increased chromatin contacts within the region [59]. TAD size was not significantly different between samples (Additional file 1: Supplementary fig 4a), and 72.5% of TADs identified in cells migrated with constriction overlapped by ≥80% with TADs in cells migrated without constriction (Additional file 1: Supplementary fig 4b). The TADs that were not conserved were significantly larger than the conserved TADs (p=2×10^−16^, R^2^=0.15, logistic regression of log transformed TAD size) (Additional file 1: Supplementary fig 4c), suggesting that the differences could be driven by boundary loss between neighboring TADs. However, cells migrated with constriction had lower contact frequencies both within and between TADs (Additional file 1: Supplementary fig. 4d). Therefore, some observed boundary deletions may be due to loss of power as opposed to true changes in TAD structure. Regardless, conserved TADs were more likely to be found in compartment A than non-conserved TADs (Fig 3d), suggesting again that the A compartment is more stable after migration with constriction.

**Figure 4.**
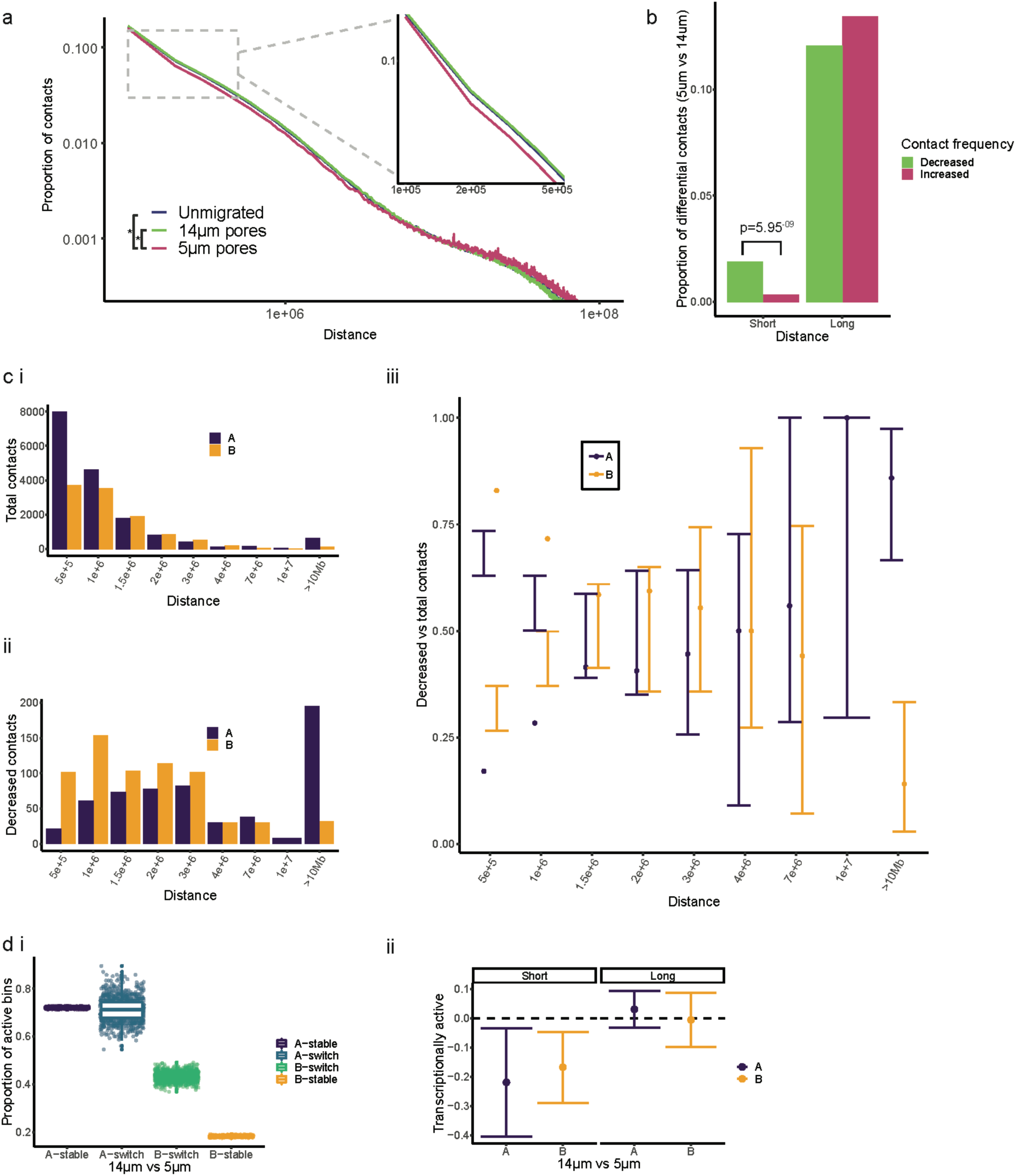
Short range contacts were depleted after migration with constriction. a) The whole genome contact matrix at 100kb resolution was normalized with iterative correction and eigenvector decomposition (ICE), and distance dependent contact probability plotted on a log-log scale. There was no significant difference between the distributions of unmigrated cells, and cells migrated without constriction (14μm pores) (KS test, p=0.14), while cells migrated with constriction (5μm pores) have a significantly different distribution from both unmigrated and migrated without constriction (KS test 5.74×10^−9^ and 3.11×10^−13^ respectively). The inset highlights the rapid decay of contact frequency between 100kb and 500kb in migration with constriction. b) Differential intrachromosomal contacts between migration with and without constriction were called at 100kb resolution, tiling across the chromosome in 40kb bins. Long range (>1Mb) differential contacts were equally likely to be lost or gained, while short range (100kb-1Mb) were significantly more likely to be lost after migration with constriction. Regions involved in a contact were assigned to compartment A or B. c) Distribution of disrupted contacts. Bin sizes are uneven. i) Number of significant contacts (FDR<0.05) in compartment A and B in cells migrated without constriction. ii) Number of significantly decreased contacts (FDR<0.1) in compartment A and B between migration with and without constriction. iii) Bootstrapping of significant contacts found a strong enrichment of decreased short range (<1Mb) contacts in compartment B. Error bars represent 99% CI of the expected proportion of contacts in compartment A or B. based on 10,000 bootstraps. d) 100kb bins were defined as transcriptionally active if they contained one of more expressed gene. i) Stable A compartment regions were most likely to be active, and stable B compartment genes were least likely to be active. ii) Bootstrapping of total contacts found that disrupted contacts in either compartment A or B were less transcriptionally active than total contacts in the same compartment.

Our results indicate that the two major features of genome organization, compartments (A and B) and topologically associated domains (TADs), were mostly conserved after migration with or without constriction. Those disruptions that did occur, occurred preferentially within the heterochromatin-associated compartment B.

### Silent, short range contacts are disrupted after migration with constriction

Regions of chromatin that are close in linear space are more likely to come into contact than those that are further apart [57], but the rate of degradation of contact frequency across distance can vary. For instance, during neutrophil differentiation chromatin undergoes supercoiling, resulting in a loss of contacts below 3 Mb apart, and an increase in long range contacts above 3 Mb [47]. We normalized pooled contact matrices (100kb resolution) with iterative correction and eigenvector decomposition (ICE), and calculated the contact frequency at each distance (Fig 4a). Cells that were unmigrated or migrated without constriction had very similar contact frequency distributions, as a function of distance (KS test, p=0.14). Cells migrated with constriction exhibited contract frequency distributions that were significantly different from both unmigrated and migrated without constriction (KS test 5.74×10^−9^ and 3.11×10^−13^ respectively).

HOMER [58] was used to identify significant intrachromosomal contacts >100kb apart (FDR<0.05) at 100kb resolution, and to compare normalized contact frequencies between migrated and unmigrated cells Short range contacts (<1Mb apart) had a significantly higher proportion of decreased than increased frequency contacts after migration with constriction compared to migration without constriction (2 sample test for equality of proportions, p=6×10^−9^, Fig 4b). In contrast, long range contacts (>1Mb apart) had no significant difference between the number of contacts lost or gained (p=0.39, Fig 4b).

Total and decreased contacts were assigned to compartment A or B and binned according to distance between contacting regions (Fig 4c). Total significant contacts identified in cells migrated without constriction were predominantly short range, and all ranges had a higher proportion of A compartment contacts (Fig 4ci). Differential contacts had a more normal distribution, and while the long range contacts were more frequently in the A compartment, short range contacts were preferentially B compartment (Fig 4cii). To test if these apparent differences were significant, we sampled with replacement from the pool of total contacts and found the proportion of A and B compartment contacts at each range. 99% confidence intervals from 10000 bootstraps show a significant enrichment of disrupted B compartment contacts <1Mb apart, compared to total contacts (Fig 4ciii). This trend was also apparent in contacts that increased in frequency between migration with and without constriction, although the small number of short range increased contacts makes the conclusion less clear (Additional file 1: Supplementary fig 5).

**Figure 5.**
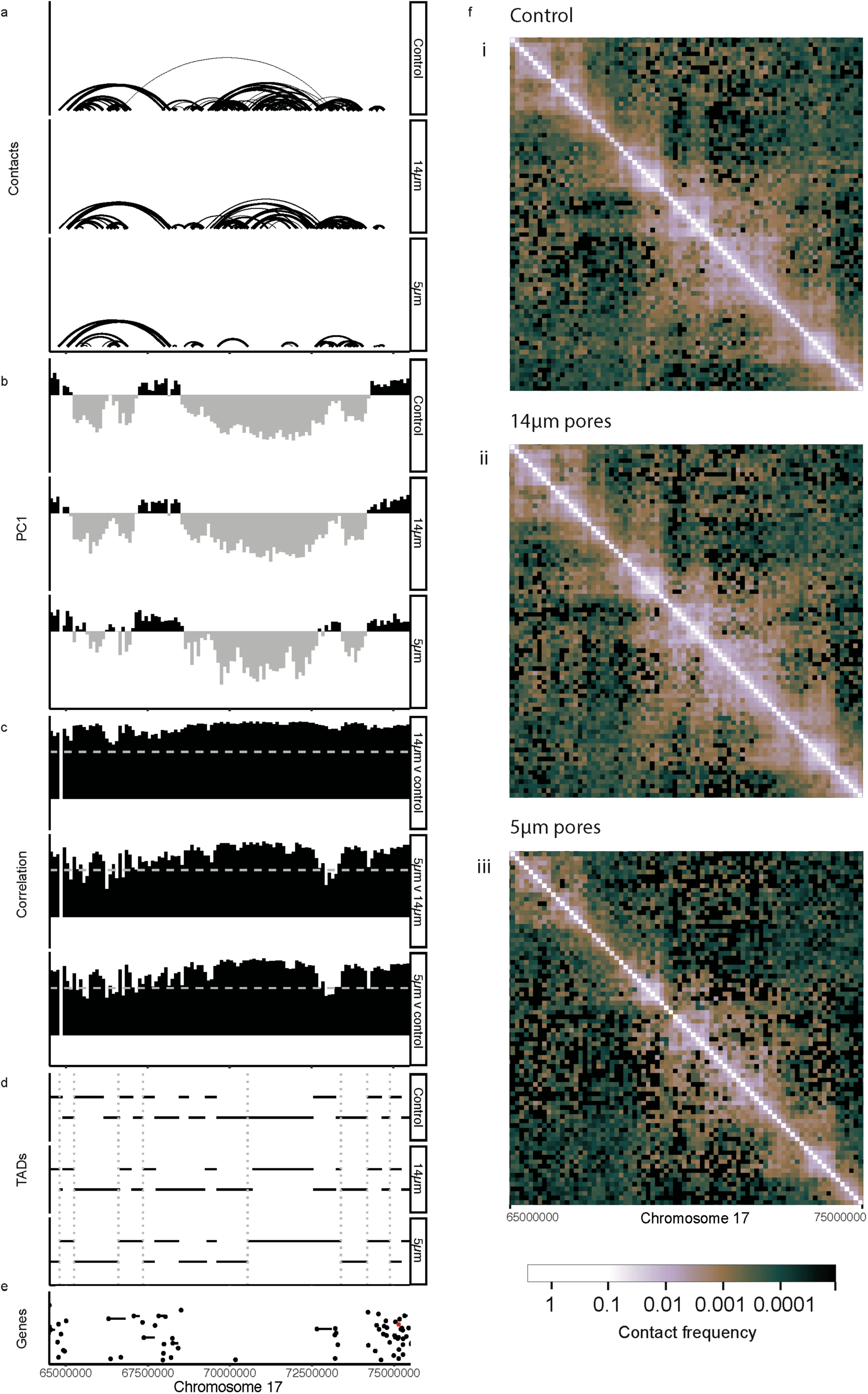
An example region of chromosome 17 showing genome organization features following migration with or without constriction. a) Significant contacts (FDR<0.05, 100kb resolution) in unmigrated cells, and cells migrated with or without constriction. Line thickness indicates significance. b) Hi-C heatmap PC1 values, indicating compartment A (positive, black) and B (negative, grey) in the three conditions. c) Correlation values show how similar the contact patterns are between two conditions. Comparisons between all three conditions are shown here, with a dotted line at R=0.6 indicating the cutoff used to call compartment switching. d) Domain (TAD) locations for all three conditions are shown here. The boundaries of TADs that are not conserved between migration with and without constriction are indicated with dashed grey lines. e) The positions of expressed genes, and log mean gene expression of the two migration conditions. One gene in this region was differentially expressed (red) between migration with and without constriction, however it was in a region with stable compartments, TADs, and contacts. f) Hi-C heatmaps of the ICE normalized contact frequency at 128kb resolution, visualised in HiGlass [107]. Lighter colour indicates higher contact frequency between regions in i) unmigrated cells, ii) cells migrated without constriction, and iii) cells migrated with constriction.

We investigated transcriptional activity in the migration dependent disrupted regions. Although the A compartment has a much higher proportion of transcriptionally active regions, with 67% of bins containing at least once transcribed gene, the B compartment still contains transcriptionally active regions (17%), and 40% of the non-conserved B regions contain actively transcribed gene(s) (Fig 4d). We identified total and decreased contacts in the A and B compartments that contained at least one expressed gene, and found that short range disrupted contacts in either compartment were less likely to be transcriptionally active (A: p=0.001, B: p=0.008). However long range contacts were equally likely to be active (A: p=0.40, B: p=0.97).

Collectively these results indicate that while differential long range (>1Mb) contacts appear to be randomly sampled from the pool of total contacts, differential short range (<1Mb) contacts are more likely to be disrupted, enriched in compartment B, and depleted for transcriptional activity.

### Transcriptionally active DNA is protected from remodeling

The disruption of inactive chromatin was observed at the contact, TAD, and compartment levels. This effect was not driven by concentrated disruption to specific regions, but instead was distributed across the genome (Additional file 1: Supplementary fig 6a-c). Although disruption to compartments, TADs, and contacts all preferentially occurred in inactive chromatin, the location of disruptions were not always consistent. For example, this region (17q24.1-q25.1) on chromosome 17 shows compartment disruption and TAD merging, but there are no significantly down-regulated short range contacts (Fig 5).

Collectively our results show that inactive chromatin organization was disturbed in cells migrated with constriction, as demonstrated by disruption to TADs and short range contacts. This disruption also affected contact patterns as detected by PCA analysis, and was not associated with transcriptional activity. This is consistent with transcriptionally active chromatin being protected from disruption during migration.

## Discussion

Our simplified model of migration was able to isolate the effects of neutrophil-like cell passage through the migration chamber with and without constriction. The induction of inflammatory, chemotaxis, and proliferative genes in cells that had migrated through large pores supports both the validity of our migration model, and roles for crowding [60] and fluid shear [61] in neutrophil activation.

Transcriptional changes specifically associated with constricted migration were enriched for genes involved in energy intake, specifically the pyruvate and lactate transporters *SLC16A7* [62] and *SLC16A3* [63]. Anaerobic glycolysis, which produces ATP via the glycolysis of pyruvate to lactate, is the neutrophils’ main form of ATP generation [64]. Cytoskeleton remodeling is energy intensive, and lactate production has the inevitable side effect of acid production [65]. Milieu acidification is a characteristic of inflammatory sites [66], and may be a direct consequence of energy intensive migration. Intriguingly, metastatic cancer cells also rely on aerobic glycolysis for movement, despite also generating ATP via mitochondrial respiration [67].

Transcript levels of the Rab GTPases also changed with constricted migration-associated remodeling. The Rab GTPases are associated with vesicle mobilization [68], which is a critical part of the neutrophil immune response [69,70]. Granule exocytosis is also regulated by adherence [71,72], cytoskeleton remodeling [73], and migration [74], supporting a role of cell shape change and mechanotransduction in priming neutrophil antibacterial responses.

Recent studies have shown that heterochromatin forms a phase-separated compartment within the nucleus [56,75]. These heterochromatic phases act like viscous ‘droplets’, and therefore should remain demixed as the nucleus changes shape [76]. Our results support this model as the widespread disruption of short range contacts is not reflected in global changes to the compartmentalization of the nucleus. Instead the heterochromatin spatial organization is altered in response to nuclear shape change [25], possibly in order to prevent damage and disruption to the organization of transcriptionally active euchromatin. Under this model, the multiple lobes of the neutrophil nucleus would provide a larger surface area and therefore increased force dispersion.

Neutrophils have a very low Lamin A content [18], and therefore would be expected to undergo a lower rate of nuclear rupture, but faster death due to DNA damage, based on recent studies by Denais and Raab [77,78]. This is attributed to the increased malleability of the nucleus, as Lamin A/C contributes stiffness to the nuclear envelope [18]. However chromatin composition also contributes strongly to the mechanical properties of the nucleus [20], and the high levels of heterochromatin along with the increased force dispersion could compensate for the lack of Lamin A in the nuclear envelope. Neutrophil differentiation is associated with an increase of long range contacts (>3 Mb) in compartment B [47]. The switch between short and long range contacts is not associated with transcriptional changes, indicating a structural role [47]. Indeed, the supercoiling of heterochromatin could contribute to its force absorbing properties during migration.

There are a several features of heterochromatin that could explain its increased sensitivity to migration with constriction [20,25]. Firstly, its peripheral location and tethering to the nuclear lamina [40,79] means it cannot be isolated from any expansion or movement of the nuclear envelope. Secondly, its rigidity [21] may lead to slower recovery from disruptions. Thirdly, transcriptionally active interactions are stabilized both by direct protein-DNA interactions [80] and by phase separated sub-compartments [46,81,82], and therefore may be more resilient to nuclear remodeling. A higher frequency of these stabilized interactions could explain the reduced disruption seen in active chromatin.

Microscopy of aspirated nuclei and magnetically twisted cells has shown that chromatin linearizes as the nucleus stretches [30,31]. These studies focused on loci that were a repetitive region [30] and a bacterial artificial chromosome [31]. Therefore, it is likely that these loci were located in the heterochromatic phase. Euchromatic loci may prove to be less malleable, although we would expect to see linearization to some extent, just as we did see disruption occurring in active chromatin.

The enrichment of disrupted contacts in transcriptionally inactive loci even within compartment A leads us to believe that migration-associated chromatin remodeling is not a major contributor to gene regulation. However, we were not able to identify specific promoter-enhancer contacts due to the resolution of the Hi-C data. Promoter-capture Hi-C [83] after migration may reveal a finer level of chromatin remodeling in response to nuclear shape change. Nascent RNA-sequencing would also provide more accurate measures of transcriptional activity, as total RNA-sequencing captures the result of both RNA production and degradation [84]. Moreover, the spatial organization and transcriptome may have different characteristics at different time points after migration. The acute changes may activate pathways and feedback loops and result in stable changes in gene expression. Alternatively, continued migration may be necessary to retain the transcriptional signature of a migrated cell. The stability of the chromatin contact changes is also unknown. As such, the disruption of short range contacts observed after 30 minutes may reflect a ‘recovering’ nucleus that is converging on the unmigrated cell structure, or a ‘remodeling’ nucleus that is in the process of gaining a distinct nuclear structure.

We controlled for confounding factors by including large pore migration, and comparing transcriptional changes to those occurring during differentiation. However, it remains possible that migration selected for a subpopulation of cells, or that some effects were mediated by cell density changes and other artefacts of the migration chamber. Single cell RNA-seq is able to detect rare cell populations [85], and thus could deconvolute the effects of migration from possible changes in population structure. Although not in the scope of this study, further experiments investigating acute chromatin conformation changes in response to a variety of stimuli, such as cytokines and contact inhibition, might help to further isolate the effects of migration with constriction.

A wider range of pore sizes would also provide greater insight into the nuclear responses to acute cell shape change. Neutrophils can migrate through gaps as small as 1.5μm diameter [86], substantially smaller than the 5μm diameter pores used in this study. We would expect to see more extreme phenotypes with smaller pores, although it is unclear how the increased disruption would manifest. Given the high rate of recovery that occurs following nuclear rupture, it is plausible that TADs and compartments are either maintained or rapidly reformed [77]. Whether inactive chromatin is able to provide any protection during migration through very small pores is unknown. Further experiments could assess the chromatin content of deformation-induced nuclear blebs and protrusions [21].

All cells in the human body must make a compromise between stiffness and flexibility of the nucleus, which is controlled by the nucleoskeleton and chromatin composition[20]. Migration through a constriction is an excellent model for this compromise, as it requires resistance to mechanical forces generated by the cytoskeleton, yet the nucleus must change shape. The stiff nuclei of cancer cells rupture during migration, but lowering Lamin A levels to decrease stiffness leads to increased cell death [77,78]. Similarly, loss or mutation of Lamin A leads to muscular dystrophy and cardiac failure [4,87], as muscle nuclei are unable to survive the repeated force of contraction. However, extreme stiffness also causes cardiac dysfunction [88,89]. Lamin A is evenly distributed across the nuclear periphery and therefore nuclear lamina stiffness is consistent across the nucleus. In contrast, it is possible for chromatin to form stiff ‘foci’ for the cytoskeleton to push against [25], while remaining flexible to allow shape change. Thus, chromatin may alleviate the loss of Lamin A protection in nuclei that undergo force-induced shape change.

The mechanisms that interact to confer nuclear properties during mechanical stress are likely to be complex and partially redundant. For instance, knockdown of SUN1, a semi-redundant component of the LINC complex [90], rescues Lamin A knockout mice from Emery-Dreifuss muscular dystrophy [91]. Notably, granulocytic HL-60/S4 cells have low levels of SUN1 [92], which may contribute to their enhanced survival compared to Lamin A knockout cells [77,78]. Assessing nuclear rupture and DNA damage following very small pore migration in cancer cells derived from rigid, non-motile tissues [77,78,93,94] is required for our understanding of metastasis and cancer evolution. However, investigating chromatin responses in nuclei adapted to migration is also important for our understanding of non-pathogenic cell processes.

Taking an integrative approach to studying nuclear mechanics is not trivial. However, understanding the interplay between Lamin A, heterochromatin, SUN1, and other nuclear components remains an important frontier in cells undergoing migration and contraction during neuronal development, bone healing, cardiovascular function, skeletal muscle growth, and the immune response.

## Conclusion

We have shown that constricted migration disproportionally disrupts inactive chromatin, possibly protecting transcriptionally active chromatin and minimizing transcriptional dysregulation during migration through a 5μm pore. Our study contributes to the growing understanding of heterochromatin as a key player in the structure and mechanosensitivity of the nucleus.

## Methods

### Cell culture

HL-60/S4 cells (available from ATCC #CCL-3306) were cultured at 37°C, 5% CO_2_ in RPMI 1640 (ThermoFisher) supplemented with 10% fetal bovine serum (Moregate Biotech), 1% penicillin/streptomycin (ThermoFisher), and 1% GlutaMAX™ (ThermoFisher). Cells were differentiated into a granulocytic phenotype with 1μM all trans retinoic acid (ATRA) dissolved in ethanol (Sigma Aldrich) for four days as previously described [51,95]. To visualize nuclear morphology, cells were cytospun at 500 rpm for 5 minutes onto slides, then stained with Wright-Giemsa (Sigma-Aldrich).

### Flow Cytometry

Flow cytometry analysis of the cell surface antigen, CD11b, was performed as described previously [18]. Briefly, 10^6^ cells were washed in PBS, blocked with human IgG (Sigma-Aldrich), then incubated with Alexa Fluor 700 Mouse Anti-Human-CD11b for 30 minutes. The proportion of live/dead/apoptotic cells was assessed with the Alexa Fluor® 488 annexin V/Dead Cell Apoptosis Kit with Alexa Fluor® 488 annexin V and PI (ThermoFisher). Briefly, 5×10^5^ cells were washed and incubated with Alexa Fluor® 488 annexin V and 100 μg/mL PI for 15 minutes. Fluorescence was measured with a FACSAria II SORP cell sorter (BD Biosciences). Analysis was performed in FlowJo v10.

### Migration

A Boyden chamber (NeuroProbe, BY312) was modified by cutting the chamber in half, leaving the upper and lower wells separated by a filter held in place by an acetal retainer. A trimmed p200 pipette tip was inserted in the opening of the lower well, and a hole was drilled on the opposite side of the lower well so a tube could be connected to a syringe. These modifications facilitated continuous flow of media through the lower well (Fig 1a). Migration assays were carried out in a hypoxic glove box (Coy Laboratory Products) at 37°C, 5% CO_2_, 20% O_2._ Media was perfused at 0.2mL/min with a syringe pump (New Era Pump Systems, NE-1002X). Growth media was used in both upper and lower wells. Polycarbonate membranes with 5μm diameter pores or 14μm diameter pores (NeuroProbe) were imaged using scanning electron microscopy. Migrated cells were collected and processed for sequencing every 30 minutes for 3-4 hours (5μm pores) or 1 hour (14μm pores). Unmigrated control cells were not exposed to the migration chamber, and were processed for sequencing during the first 30 minutes of the migration assay. All processed time points were pooled and frozen at -80°C prior to RNA extraction or Hi-C library preparation.

### RNA sequencing

Cells were lysed in TRiZOL LS (Invitrogen) and phase separated with chloroform. At this point four 5μm migration assays were pooled per replicate in order to extract enough RNA for sequencing. RNA was extracted with the Qiagen RNeasy micro kit (Qiagen, 74004). RNA and lncRNA libraries were prepared by Annoroad Gene Technology Co., Ltd. (Beijing, China) using ribosomal RNA depletion with RiboZero Magnetic Gold Kit (Human/Mouse/Rat) and sequenced on the Illumina Hi-seq X 150PE.

### Hi-C

Whole genome chromosome conformation capture (Hi-C) was performed as in [96] with restriction enzyme *Mbo*I (NEB).[96] Formaldehyde fixed cells from 4-5 migration assays were pooled to reach 2 million cells required. The library was prepared for Illumina sequencing with the TruSeq Nano kit (Truseq Nano DNA LT Sample Preparation Kit). One library was sequenced per lane of the Illumina HiSeq X 150PE.

### RNA-seq analysis

Read quality was confirmed with FastQC v0.11.4. Reads were aligned to hg38 and gencode annotations v27 using STAR v2.5.3a with default settings. FeatureCounts v1.5.2 was used to aggregate transcripts for gene-level analysis and quantify the reads with GENCODE annotations v27 (Additional file 3: Supplementary table 7). MultiQC was used to summarise FastQC, STAR, and FeatureCounts outputs [97]. Expressed genes were filtered using default settings in DEseq2 v1.16.1. PCA of variance stabilized transcripts confirmed clustering by condition (Additional file 1: Supplementary fig 3a). Differentially expressed genes were identified at FDR<0.05 using DEseq2 v1.16.1. Gene ontology enrichments were calculated with TopGo v2.28.0 using the weight01 algorithm and the Fisher statistic. Categories containing <2 genes were filtered out, and p values adjusted for FDR.

### Hi-C analysis

Read quality was confirmed with FastQC v0.11.4. Reads were aligned to hg38 and filtered using HiCUP v0.5.9 (Additional file 3: Supplementary table 8). MultiQC was used to summarize FastQC and HiCUP outputs [97]. High correlation between replicates was confirmed (R>0.994, Additional file 3: Supplementary table 9), then libraries were pooled and downsampled so that the three conditions (control, 5μm pores, 14μm pores) had the same number of valid reads (144,382,298). HiCUP output was converted to HOMER [58] input using the hicuptohomer script provided by HiCUP. Compartments were identified by performing PCA on each on each chromosome in HOMER v4.9.1 at 100kb resolution on a matrix normalised with the default HOMER method. TSS locations were used to seed active regions to determine the correct sign of the PC1 value. Chromosomes were excluded from compartment analysis if the first principal component corresponded to the chromosomal arms (chromosomes 4,5,19,21,X). Correlation matrices were compared to identify compartment switching with the getHiCcorrDiff.pl script provided by HOMER. Significant and differential intrachromosomal contacts were identified in HOMER at 100 kb resolution. The Hi-C matrices were normalized using the default HOMER method. Reads supporting contacts were counted in 100kb windows, tiling across the chromosome in 40kb bins to avoid penalizing boundaries. This is known in HOMER as 40kb resolution, 100kb super-resolution. HOMER was also used to generate raw chromosomal matrices at 50 kb resolution for topologically associated domain (TAD) analysis in TopDom v0.0.2 using a window size of 5, as performed in [98]. Iterative correction and eigenvector decomposition (ICE) was performed with cooler v0.7.1 [99]. Hi-C heatmaps were generated with HiGlass v1.1.5 [100]. Genomic interval arithmetic was performed with bedtools v2.27.0 [101] and plyranges v1.0.3 [102]. Plots, excluding heatmaps, were generated in R 3.5.1 [103] with dplyr [104] and ggplot2 [105]. Color Oracle (colororacle.org) was used to assist color choices in all figures [106].

## Declarations

### Ethics approval and consent to participate

Not Applicable

### Consent for publication

Not Applicable

### Availability of data and materials

The datasets generated and analyzed during the current study are available in the GEO repository, accession number GSE115634. Scripts and data to reproduce analysis of processed data is available on github url: https://github.com/jacel/migration-chromatin. Additional datasets required for the analyses are available on Figshare: 10.17608/k6.auckland.6943112. Data from [95] is available from the NCBI Short Read Archive http://www.ncbi.nlm.nih.gov/bioproject/303179 [55].

## Authors’ Contributions

All authors contributed to conception and design of the study. EJ performed the experiments, analyzed and interpreted the data, and wrote the manuscript with support from JMO, JKP, and MHV. All authors contributed to revisions and approved the final manuscript.

## Acknowledgements

The authors would like to thank Peter Shepherd and the members of the JMO lab group for comments and discussion. We thank Stephen Olding in the Auckland Bioengineering Institute for assistance with modifying the migration chamber, Jacqueline Ross in the Biomedical Imaging Research Unit, University of Auckland for assistance with developing the assay, and Stephen Edgar in the Core Flow Cytometry Facility, FMHS, University of Auckland for assistance with flow cytometry.

## Additional files

### Additional File 1: Figures 1-4. 1

HL60 cells differentiated into neutrophil-like cells and migrated through two pore sizes. **2.** SEM of porous membranes. **3.** RNA-seq summary plots, and comparison of migration with differentiation. **4.** Topological domain changes after migration through two pore sizes. **5.** Compartment status of increased frequency contacts between migration with and without constriction. **6.** Genome wide distribution of disrupted contacts, compartments, and TADs.

### Additional File 2: Table 1

*Significantly differentially expressed genes*. All genes with significantly differential gene expression (FDR<0.05) between either 5μm pores compared to control, 14μm pores compared to control, or 5μm pores compared to 14μm pores. Sets refer to those described on page 4 (*Transcriptional changes after migration and remodeling are distinct*). HUGO gene names, Ensembl gene names and IDs are provided as identifiers. Columns suffixed with ‘_5’ represent results from 5μm pore compared to control. Columns suffixed with ‘_14’ represent results from 14μm pore compared to control. Columns suffixed with ‘_5v14’ represent results from 5μm pore compared to 14μm pore. The ‘padj’ shows the FDR adjusted p-value.

## Additional File 3: Table 2-5, 7-9. 2

Gene ontology enrichment (Biological process) of Fig 2B. **3.** Gene ontology enrichment (Biological process) of Fig 2C. **4.** Gene ontology enrichment (Biological process) of Fig 2D. **5.** Gene ontology enrichment (Molecular function) of Fig 2D. **7.** RNA-seq QC. **8.** HiCUP QC. **9.** Hi-C replicate correlations.

## Additional File 4: Table 6

*PC1 values and correlation between the interaction patterns of 100kb bins.* Each 100kb bin has both a PC1 value indicating compartment status, and an interaction pattern with every other bin in the chromosome. When bins have a low correlation (R<0.6) and an opposite PC1 value, this is considered a disruption to the compartment status of this region. Calculated using the runHiCpca.pl and getHiCcorrDiff.pl scripts in HOMER. Regions with missing values, and chromosomes with PC1 values indicating chromosome arm, are excluded from this table.

